# Continuous theta burst TMS of area MT+ impairs attentive motion tracking

**DOI:** 10.1101/601807

**Authors:** Arijit Chakraborty, Tiffany T. Tran, Andrew E. Silva, Deborah Giaschi, Benjamin Thompson

**Affiliations:** School of Optometry and Vision Science, University of Waterloo; Department of Ophthalmology and Visual Sciences, The University of British Columbia; Chicago College of Optometry, Midwestern University; Department of Psychology, University of California Los Angeles

**Keywords:** multiple object tracking, transcranial magnetic stimulation, higher visual processing, extrastriate visual cortex, covert attention

## Abstract

Attentive motion tracking deficits, measured using multiple object tracking (MOT) tasks, have been identified in a number of visual and neurodevelopmental disorders such as amblyopia and autism. These deficits are often attributed to the abnormal development of high-level attentional networks. However, neuroimaging evidence from amblyopia suggests that reduced MOT performance can be explained by impaired function in motion sensitive area MT+ alone. To test the hypothesis that MT+ plays an important role in MOT, we assessed whether modulation of MT+ activity using continuous theta burst stimulation (cTBS) influenced MOT performance in participants with normal vision. An additional experiment involving numerosity judgements of MOT stimulus elements was conducted to control for non-specific effects of MT+ cTBS on psychophysical task performance. The MOT stimulus consisted of 4 target and 4 distractor dots and was presented at 10° eccentricity in the right or left hemifield. Functional MRI-guided cTBS was applied to left MT+. Participants (n = 13, age:27 ± 3) attended separate active and sham cTBS sessions where the MOT task was completed before, 5 mins post and 30 mins post cTBS. Active cTBS significantly impaired MOT task accuracy relative to baseline for the right (stimulated) hemifield 5 mins (10 ± 2% reduction; t_12_ = 1.95, p = 0.03) and 30 mins (14 ± 3% reduction; t_12_ = 2.96, p = 0.01) post stimulation. No impairment occurred within the left (control) hemifield after active cTBS or for either hemifield after sham cTBS. Numerosity task performance was unaffected by cTBS. These results highlight the importance of lower-level motion processing for MOT and suggest that abnormal function of MT+ alone is sufficient to cause a deficit in MOT task performance.

## Introduction

Attentive motion tracking is a fundamental component of vision that supports navigation and interpretation of dynamic visual environments. Multiple object tracking (MOT) can be used as an experimental task to measure attentive motion tracking (Alvarez & Franconeri, 2007; Cavanagh & Alvarez, 2005; C. S. Ho et al., 2006; Meyerhoff, Papenmeier, & Huff, 2017; Pylyshyn, 2004; Pylyshyn & Storm, 1988). A typical MOT task requires the use of covert visual attention to track visual targets presented within an array of distractors. Targets are cued at the start of each trial and both targets and distractors move along unique trajectories during the trial. Targets are discriminated from distractors using full or partial report at the end of a trial. Adults with normal vision can covertly track four to five targets (Pylyshyn, 2004).

MOT tasks activate a distributed network of neural areas in humans including the frontal eye fields (FEF), anterior intraparietal sulcus (AIPS), posterior intraparietal sulcus (PIPS), superior parietal lobule (SPL), precuneus and motion sensitive area MT+ (Culham et al., 1998; Culham, Cavanagh, & Kanwisher, 2001; Howe, Horowitz, Morocz, Wolfe, & Livingstone, 2009; Kastner & Ungerleider, 2000; Merkel, Hopf, Heinze, & Schoenfeld, 2015; Ungerleider, 2000). Activity within the FEF and SPL may be due to involuntary planning of saccadic eye movements (SPL) that subsequently need to be suppressed (FEF) during an MOT task (Burman & Bruce, 2009; Doricchi et al., 1997; Howe et al., 2009; Priori, Bertolasi, Rothwell, Day, & Marsden, 1993). Control of covert attention is likely to involve the AIPS and PIPS. Activity in both areas is associated with attention to moving targets (Donner et al., 2000; Howe et al., 2009). In addition to evidence from fMRI demonstrating a correlation between AIPS activity and MOT, a role for the AIPS in MOT has been supported by non-invasive brain stimulation studies. Specifically, inhibitory repetitive transcranial magnetic stimulation (rTMS) of the intraparietal sulcus (IPS) impairs MOT (Battelli, Alvarez, Carlson, & Pascual-Leone, 2009) and excitatory anodal transcranial direct current stimulation (tDCS) of the AIPS improves MOT (Blumberg, Peterson, & Parasuraman, 2015).

The role of area MT+ in attentive motion tracking is less clear than that of the AIPS. fMRI studies indicate that MT+ activity is strongly associated with the allocation of covert attention to moving targets (Berman & Colby, 2002; Chawla, Rees, & Friston, 1999; Treue & Maunsell, 1996) and that MT+ has strong functional connectivity to the AIPS during attentive motion tracking (Howe et al., 2009). In addition, a study has provided indirect evidence for MT+ having a role in MOT deficits (Secen, Culham, Ho, & Giaschi, 2011). Amblyopia, a developmental disorder of the visual system, impairs MOT in both the amblyopic and non-amblyopic fellow eye. Using fMRI, Secen et al. (2011) observed that MT+ showed the most robust group difference between amblyopia and binocular control groups during MOT. This result suggests that an MT+ deficit underlies the MOT impairment in amblyopia. However, Battelli et al., (2009) observed that inhibitory rTMS of MT+ in normally-sighted controls did not affect MOT, implying that MT+ does not play a direct role in MOT task performance. Battelli et al., (2009) stimulated both IPS and MT+ within the same session and it is possible that a large effect of IPS stimulation may have masked any effect of MT+ stimulation. The aim of this study was to further explore the role of MT+ in attentive motion tracking using continuous thetaburst stimulation (cTBS; an inhibitory rTMS technique) (Huang, Edwards, Rounis, Bhatia, & Rothwell, 2005) and a study design that allowed for both visual hemifield and sham stimulation control conditions. cTBS reduces cortical excitability by mimicking theta-gamma oscillation coupling and impairs global motion perception in the contralateral hemifield when applied over MT+ (Cai, Chen, Zhou, Thompson, & Fang, 2014; Chen, Cai, Zhou, Thompson, & Fang, 2016; Silvanto, Muggleton, Cowey, & Walsh, 2007). Given the evidence for abnormal MT+ function in amblyopia (Bonhomme et al., 2006; Ho & Giaschi, 2009; Thompson, Villeneuve, Casanova, & Hess, 2012) and the link between the MOT impairment in amblyopia and abnormal MT+ activity (Secen et al., 2011), our hypothesis was that inhibitory unilateral rTMS of MT+ would impair MOT for stimuli presented in the contralateral visual hemifield.

The dorsal stream vulnerability hypothesis proposes that dorsal visual cortical stream areas are more susceptible to abnormal development than areas in the ventral processing stream (Braddick, Atkinson, & Wattam-Bell, 2003; Grinter, Maybery, & Badcock, 2010). Area MT+ has a central role in this hypothesis because the vast majority of relevant studies assess dorsal stream function with global motion stimuli that are specifically designed to target MT+ (Braddick et al., 2001; Kaderali, Kim, Reynaud, & Mullen, 2015; Newsome & Pare, 1988; Newsome, Britten, & Movshon, 1989; Simoncelli & Heeger, 1998). This approach is based on the idea that area MT+ is a dorsal stream area, although it has been argued that the division of processing into dorsal and ventral streams takes place after MT+ (Gilaie-Dotan, 2016; Milner & Goodale, 1995; Schenk, Mai, Ditterich, & Zihl, 2000; Schenk & McIntosh, 2010). Impaired global motion perception linked to dorsal stream vulnerability has been reported in a range of neurodevelopmental disorders including William’s syndrome (Atkinson et al., 1997, 2003; Atkinson et al., 2006; Palomares & Shannon, 2013; Reiss, Hoffman, & Landau, 2005), preterm birth (Guzzetta et al., 2009; Taylor, Jakobson, Maurer, & Lewis, 2009), autism (Brieber et al., 2010; Spencer et al., 2000), dyslexia (Edwards et al., 2004; Talcott, Hansen, Assoku, & Stein, 2000), and fetal alcohol syndrome (Gummel, Ygge, Benassi, & Bolzani, 2012). Attentive motion tracking is also impaired in conditions that affect neurodevelopment such as Down’s syndrome (Brodeur, Trick, Flores, Marr, & Burack, 2013), Turner’s syndrome (Beaton et al., 2010), chromosomal abnormalities (Cabaral, Beaton, Stoddard, & Simon, 2012), autism (Koldewyn, Weigelt, Kanwisher, & Jiang, 2013), and William’s syndrome (O’Hearn, Hoffman, & Landau, 2010; O’Hearn, Landau, & Hoffman, 2005). A common role for MT+ in global motion perception and MOT may partially explain this pattern of results. In summary, the question of whether MT+ is involved in attentive motion tracking is important for at least two reasons: 1) understanding the MOT deficits in amblyopia, and 2) understanding the effects of abnormal neurodevelopment on vision.

## METHODS

This study employed a within-subjects, single masked, sham-controlled design. An additional within-session control of stimulated vs. unstimulated hemifield was also included. Furthermore, in a separate session, a sub-set of six participants completed a control task before and after cTBS that involved discrimination of static MOT stimuli in both the stimulated and control hemifields. This control experiment was conducted to evaluate the possibility of impaired overall psychophysical task performance induced by cTBS, that was not specific to attentive tracking.

### Participants

Fifteen participants with a mean age of 27 ± 3 years (7 female) were recruited from the University of Waterloo. All participants had normal or corrected to normal vision (monocular visual acuity better than 0.00 logMAR in each eye) measured using the Freiburg Visual Acuity and Contrast Sensitivity Test (Bach, 1996, 2007). All procedures were approved by the University of Waterloo Institutional Review Board, and participants provided written, informed consent. No participants reported neurological or psychiatric conditions. The study adhered to the principles of the Declaration of Helsinki.

### Procedure

Before the TMS sessions, the participants underwent structural and functional MRI (to localize MT+) on a separate day at Robarts Research Institute, Western University. The TMS sessions were divided across 3 days. Day-1 involved active motor thresholding (AMT). Day-2 and -3 involved active/sham cTBS in a random order across participants. Day-2 began with speed threshold measurement. The same speed threshold was used for Day-3. A subset of six participants completed an additional two sessions of cTBS (active / sham) for the control experiment.

### Functional MRI localization of middle temporal cortex

Area MT+ was localized in each hemisphere for every participant using fMRI (7T Siemens Magnetom system equipped with a 12-channel, occipital-only surface coil). The imaging session began with acquisition of a T1-weighted MP2RAGE (0.7mm isotropic voxels, sagittal orientation, TR = 6000ms, TE = 2.73ms, FOV = 240mm, number of slices = 224). Two hundred and fifty functional BOLD volumes (60 slices per volume, 1×1×1.5mm voxels, TE = 19.6ms, TR = 1600ms, FOV = 208mm, flip angle = 45°, coronal orientation with outermost slice positioned on the occipital pole) were then acquired while participants viewed a visual stimulus comprised of 12 blocks of white (67 cd/m^2^) radially moving dots (400 dots, 6 deg/s) presented on a dark (4 cd/m^2^) background. Within each 16 s block, dots radially moved in and out with motion direction reversing every 1.6s. A baseline block (16s) consisting of stationary dots was presented after every motion block. The fixation point was randomly switched from “X” to “O” or “O” to “X”, and participants had to press a key whenever a change occurred. The BOLD data were preprocessed and analyzed in BrainVoyager QX (Brain Innovation, Maastricht, Netherlands). The data were preprocessed with slice scan time correction (cubic spline interpolation), 3D motion correction (trilinear/sinc interpolation), and temporal high-pass filtering. Voxels that were activated significantly more strongly by radial motion than by stationary dots were identified using a general linear model that included z-transformed head motion nuisance regressors. The cluster of significant voxels located in the expected region for left hMT+ was identified (FDR corrected q < 0.01), and the MNI coordinates of the peak were used to guide TMS coil position using BrainSight software (Rogue Research Inc., Montreal, Canada) (Figure 1).

**Figure 1:**
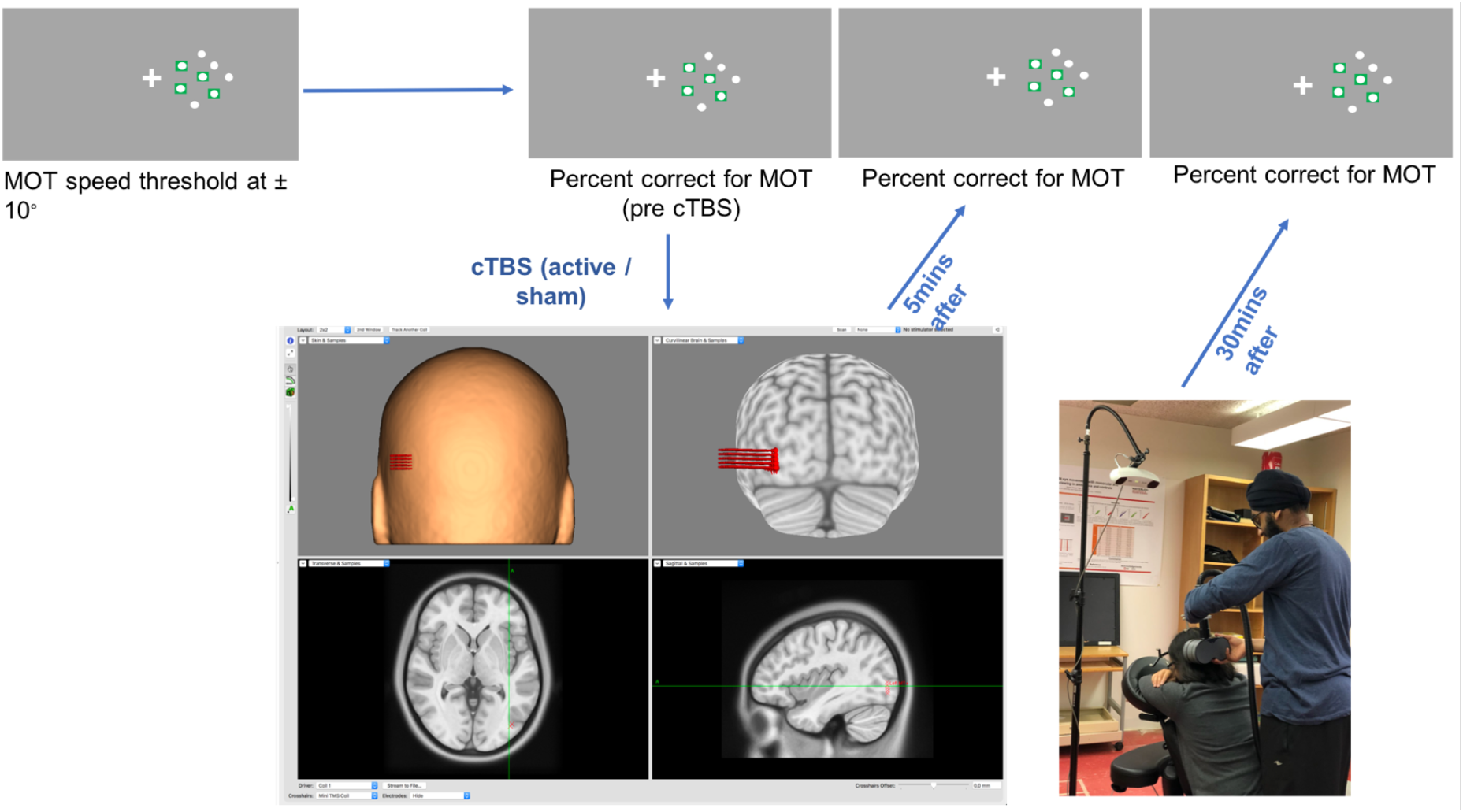
Speed thresholds for the MOT task were first measured at 10° eccentricity for both the left and right hemifields. Dot speed was fixed at threshold and percent correct scores were calculated across 40 trials before, 5mins after and, 30mins after continuous theta burst stimulation (cTBS). Neuronavigated cTBS was applied to left MT.

### Multiple-object tracking stimuli

The multiple-object tracking task, consisted of 8 white dots (diameter 1°, 4 targets and 4 distractors) on a grey background (luminance = 45 cd/m^2^) within a square aperture (14 x 14°) and was presented on a 1080p Asus (Model: PG278QR) gaming monitor. Stimuli were generated using MATLAB 2016a (MathWorks, Inc.) and Psychtoolbox 3 (Pelli, 1997). The stimuli were centered 10° to the left or right of a bright fixation cross; this eccentricity was chosen to stimulate the contralateral MT+ (Cai, Chen, Zhou, Thompson, & Fang, 2014; Chen, Cai, Zhou, Thompson, & Fang, 2016). Minimum dot spacing was maintained at 1.5° to reduce crowding effects, based on our previous work (Chakraborty, Hua, Chan, Giaschi, & Thompson, 2017). Before the dots began moving, target dots were highlighted with green square boxes at the beginning of the task for 1.5 seconds (Figure 2), following which the boxes disappeared and the 8 dots moved randomly for 5 seconds. Once the dots stopped, one of the dots was then highlighted in yellow and the observer had to report whether the highlighted dot was a target or distractor using a button press within a two-alternative force-choice design.

**Figure 2:**
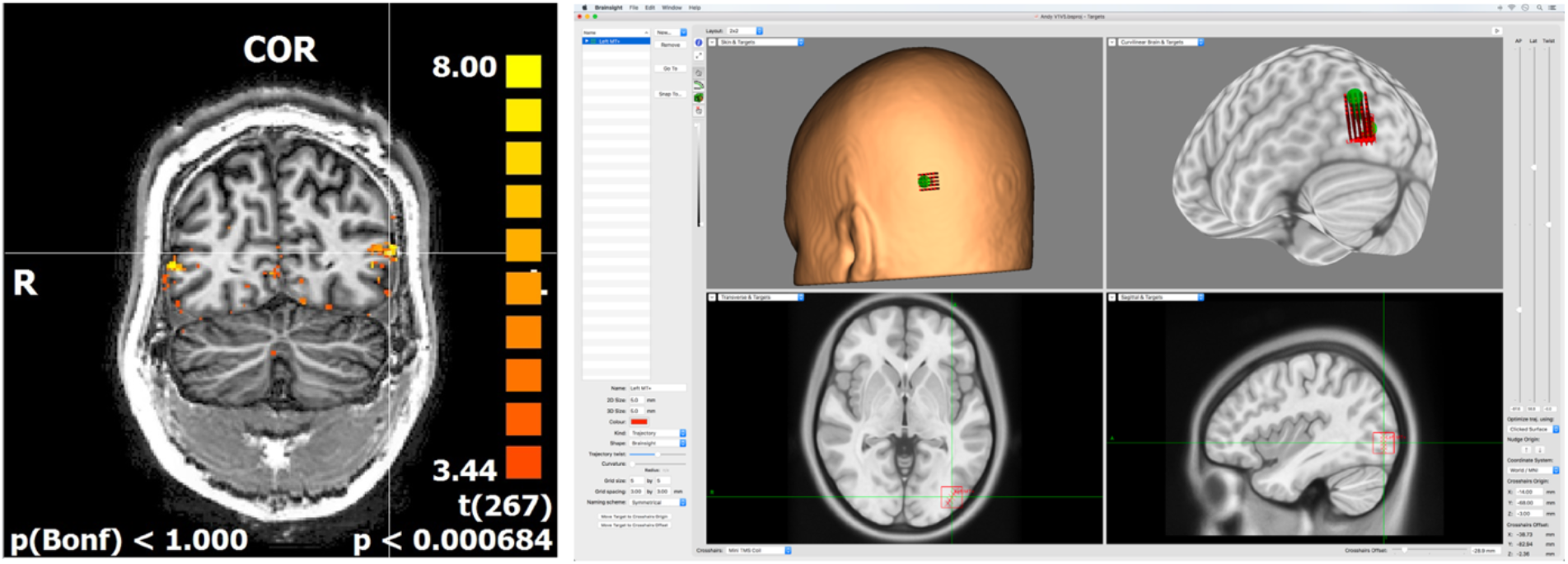
TMS stimulation sites were located using fMRI (left panel) and the Montreal Neurological Institute (MNI) coordinates of the peak intensity voxel for the left hemisphere were exported into the Brainsight neuronavigation system software (right panel). A target marker was placed on the MNI coordinate and 5×5 grid (3mm intergrid spacing) was built around the marker, which was then used to guide stimulation.

### Psychophysics

Participants viewed the MOT stimuli binocularly from 50cm with their habitual refraction correction in place. A chin rest stabilized head position. The 2-step psychophysical procedure involved 1) measurement of speed thresholds for the MOT task in each hemifield, 2) measurement of MOT task accuracy (% correct responses) with speed fixed at threshold before TMS and 5mins and 30mins post TMS.

1. A 2-up 1-down adaptive staircase was used to measure MOT speed thresholds. Baseline speed was 4°/sec. The staircase varied speed using a proportional step size of 25% until the first reversal and 12.5% thereafter. If the task could not be performed at baseline (4°/sec), baseline speed was decreased by 25%. The staircase terminated after 16 reversals and the threshold was calculated as the average of the last 12 reversals. Speed thresholds were calculated separately for each hemifield within an interleaved staircase. Participants completed a familiarization task consisting of 12 MOT trials before completing the threshold measurements. The familiarization task was similar to the test program, except that the observer tracked 2 targets out of 8 elements. In addition, 5 of the 15 participants completed an additional set of MOT speed threshold measurements while their gaze was monitored with an Eye Tribe infrared eye tracker (Copenhagen, Denmark). A 1.5 deg diameter “fixation area” was defined around the fixation point. Observers were given visual feedback (red flashing light at the fixation area) and the trial was repeated whenever the gaze shifted out of the fixation area.
2. MOT task accuracy was measured across 40 trials per hemifield with speed fixed at each individual participant’s speed threshold for each hemifield. Blocks of 10 trials were presented alternately to the right and left hemifields. The starting hemifield was randomized across participants. Task accuracy was calculated separately for each hemifield. Within each of the active and sham cTBS sessions, MOT task accuracy was measured directly before, 5 min after and 30 mins after cTBS.

### Numerosity judgement

The numerosity judgment task was conducted with static frames of the multiple object tracking stimulus. The reference and test presentations (duration: 1 sec) had 8 and 8 ± 1 elements, respectively, with an interstimulus interval of 0.5 sec. A sub-set of six observers reported which of the 2 frames had more elements. The numerosity judgement was tested at 10° eccentricity with element spacing fixed at 1.5°. Participants completed 50 trials per hemisphere (in randomly sequenced blocks of 25 trials per hemisphere) at baseline, 5 and 30 mins after active and sham cTBS delivered in separate sessions.

### Transcranial magnetic stimulation

TMS was delivered using a MagVenture MagPro X100 (MagVenture Farum, Denmark) stimulator guided by the Brainsight frameless stereotaxy neuronavigation system (Rogue Research Inc., Montreal, Canada). Active motor thresholding was conducted with a MCF B-65 coil. Active / placebo cTBS was delivered using an active / placebo B-65 coil. The coil allows for masked active / placebo stimulation as it is two-sided with an active side and a magnetically-shielded placebo side. A gyroscope in the coil provides a signal to the stimulator and the experimenter is cued to keep the coil orientation constant or to flip the coil depending on whether active or placebo stimulation is to be delivered. The maximum stimulator output of the TMS system was 159 A/μs.

### Active motor thresholding

Participants were seated in a comfortable chair. Surface electromyographic (EMG) electrodes were placed on the belly-tendon of the right first dorsal interosseus (FDI) muscle and the ground was placed on the bone of right wrist. EMG activity was monitored online using Brainsight software. Active motor threshold (AMT) was measured by asking the participant to steadily press their thumb against the arm of their chair to generate a steady motor evoked potential (MEP) of 100 μV (Deblieck, Thompson, Iacoboni, & Wu, 2008). Single pulse TMS was delivered at 40% maximum stimulator output (MSO) with the coil handle positioned at 45° from the sagittal plane (Richter, Neumann, Oung, Schweikard, & Trillenberg, 2013). The TMS coil was moved across a grid (1×1 cm^2^ spacing) covering the left motor cortex that was drawn on a template brain within the Brainsight software. Single pulses were delivered at each grid junction and also in between junctions. The region that generated the maximum EMG amplitude was identified as the “hot spot”. TMS intensity was varied in 1% MSO steps until a MEP with an amplitude greater than 200 μV (peak to peak) was produced for 5 of 10 pulses.

### Continuous theta burst transcranial magnetic stimulation (cTBS)

cTBS was administered at 100% of AMT over left MT+. Montreal Neurological Institute (MNI) coordinates of the peak left MT voxel from fMRI were used within Brain Voyager to guide coil placement on a participant-by-participant basis (Figure 1). cTBS was delivered in three pulses at 50 Hz repeated every 200ms for 40s, resulting in a total of 600 pulses. Participants sat quietly for 5 mins after cTBS.

## RESULTS

All fifteen participants exhibited strong bilateral activation within the middle temporal cortex using our fMRI protocol. Average MNI coordinates for left MT+ were X: −45 ± 5.72, Y: −69 ± 4.16, Z: −3 ± 7.56. The mean AMT was 46 ± 4.7% of maximum stimulator output. The mean speed thresholds for MOT accuracy did not vary significantly between the right (7.63 ± 2.16 deg/s) and left (8.17 ± 2.80 deg/s) hemifield (t_28_ = −0.52, p = 0.602). The five participants who were retested on the gaze-monitored MOT task during the baseline measurement had similar speed thresholds with and without eye tracking (t_4_ = −0.36, p = 0.248) (right hemifield with eye tracking: 7.41 ± 1.38 deg/s and left hemifield with eye tracking: 8.02 ± 1.11 deg/s). Two participants were excluded from the study prior to cTBS administration because their baseline MOT task accuracy was at chance (≤ 50% correct) during their first cTBS session. For the remaining 13 participants, baseline MOT task accuracy did not vary between the right and left hemifields during the active (right: 75 ± 7 % correct, left: 78 ± 10 %; t_24_ = −0.76, p = 0.453) and sham (right: 74 ± 5 %, left: 75 ± 7 %; t_24_ = −0.15, p = 0.884) cTBS sessions.

A repeated measures general linear model was conducted to test for differences in MOT task accuracy as a function of Stimulation Type (active / sham cTBS), Hemifield (left / right) and Time (baseline / 5min post cTBS / 30min post cTBS) (Figure 3). There was a significant interaction between Stimulation Type and Time (F_2,18_ = 7.18, p = 0.005). No other interactions were significant. For active cTBS, there was a significant reduction in accuracy from baseline for the right (stimulated) hemifield after 5min (10 ± 2% reduction; t_12_ = 1.95, p = 0.03) and after 30min (14 ± 3% reduction; t_12_ = 2.96, p = 0.01). The left (control) hemifield exhibited improved accuracy 30min after active cTBS (7 ± 1.5% improvement, t_12_ =−2.24, p = 0.02). For sham cTBS, accuracy improved in both hemifields equally after 30mins (right: 9 ± 2% improvement; t_12_ = −2.94, p = 0.02 and left: 8 ± 1.5% improvement; t_12_ = 1.95, p = 0.04).

**Figure 3:**
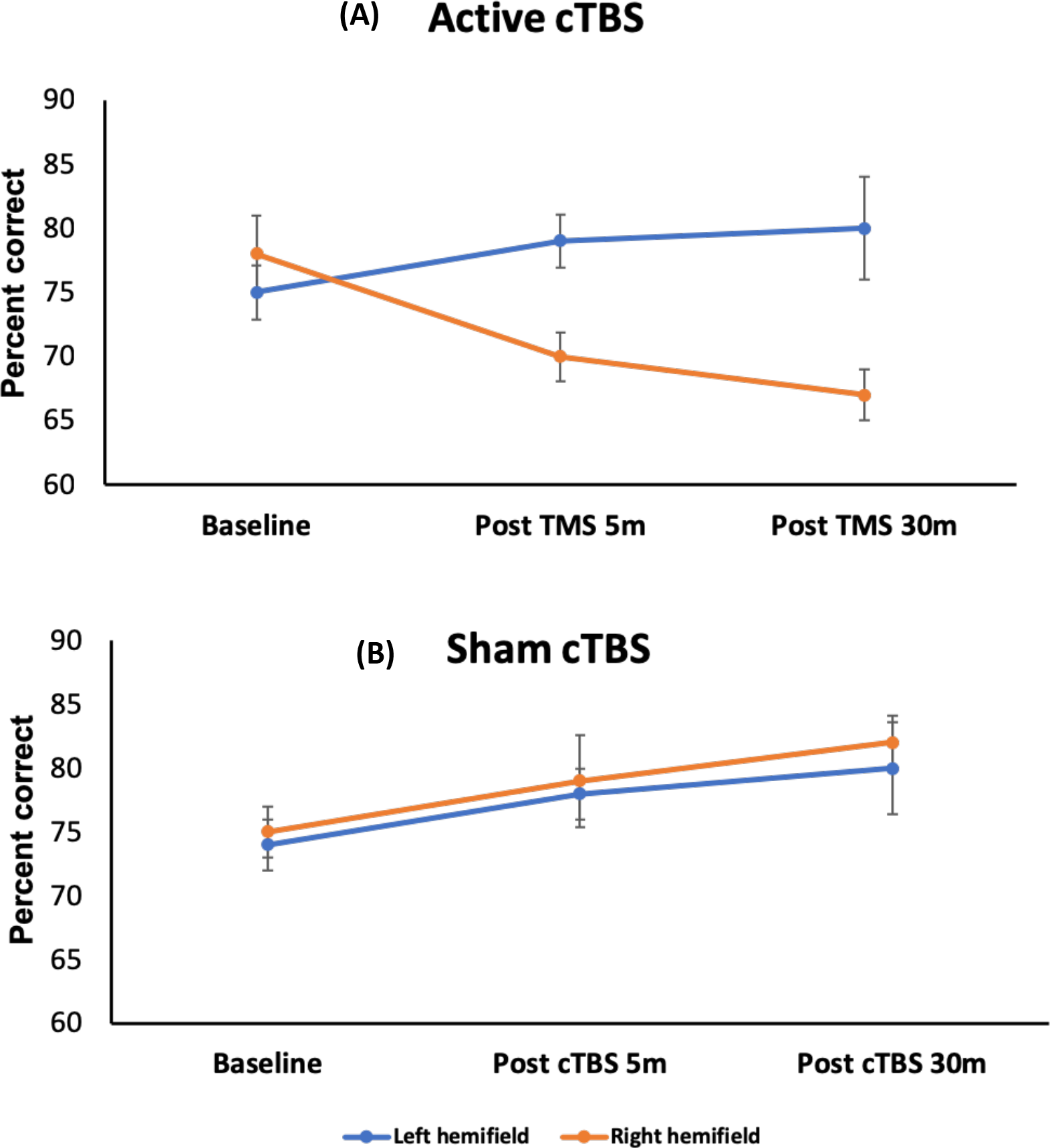
Percent correct MOT task accuracy following (A) active and (B) sham cTBS. The blue and red and green lines indicate left and right hemifield task performance, respectively, at baseline, 5min post cTBS and 30-min post cTBS. Error bars show +/− 1 SEM.

Response accuracy for the control numerosity judgement task increased 5mins and 30mins after active cTBS in both stimulated [F_2,15_ = 7.97, p = 0.004] (post-5min: 5 ± 4% increase, t_5_ = −3.50, p = 0.02; post-30min: 12 ± 6% increase, t_5_ = −5.12, p = 0.004) and control hemifields [F_2,15_ = 9.09, p = 0.002] (post-5min: 5 ± 3% reduction, t_5_ = −3.50, p = 0.02; post-30min: 11 ± 4% reduction, t_5_ = −5.83, p = 0.002). Similarly, improvement was also observed with sham cTBS in both the stimulated F_2,15_ = 7.56, p = 0.005] (post-5min: 7 ± 4% reduction, t_5_ = −4.50, p = 0.006; post-30min: 13 ± 2% reduction, t_5_ = −15.49, p < 0.001) and control hemifields F_2,15_ = 5.56, p = 0.015] (post-5min: 5 ± 4% reduction, t_5_ = −2.98, p = 0.03; post-30min: 12 ± 4% reduction, t_5_ = −7.41, p = 0.001). Finally, a general linear model was constructed to compare the effect of active cTBS on the MOT task and the numerosity task for these six participants. The ANOVA model was conducted on the percent correct scores in the right (stimulated) hemifield and had factors of Test Type (MOT vs. numerosity) and time (Pre, 5 and 30 mins post active cTBS). The effect of active cTBS differed significantly between the two tests (significant interaction between Test Type and Time, F_2,10_ = 30.74, p < 0.001; figure 4). This indicated that the effect of cTBS on MOT was not simply due to a non-specific disruption of psychophysical task performance.

**Figure 4:**
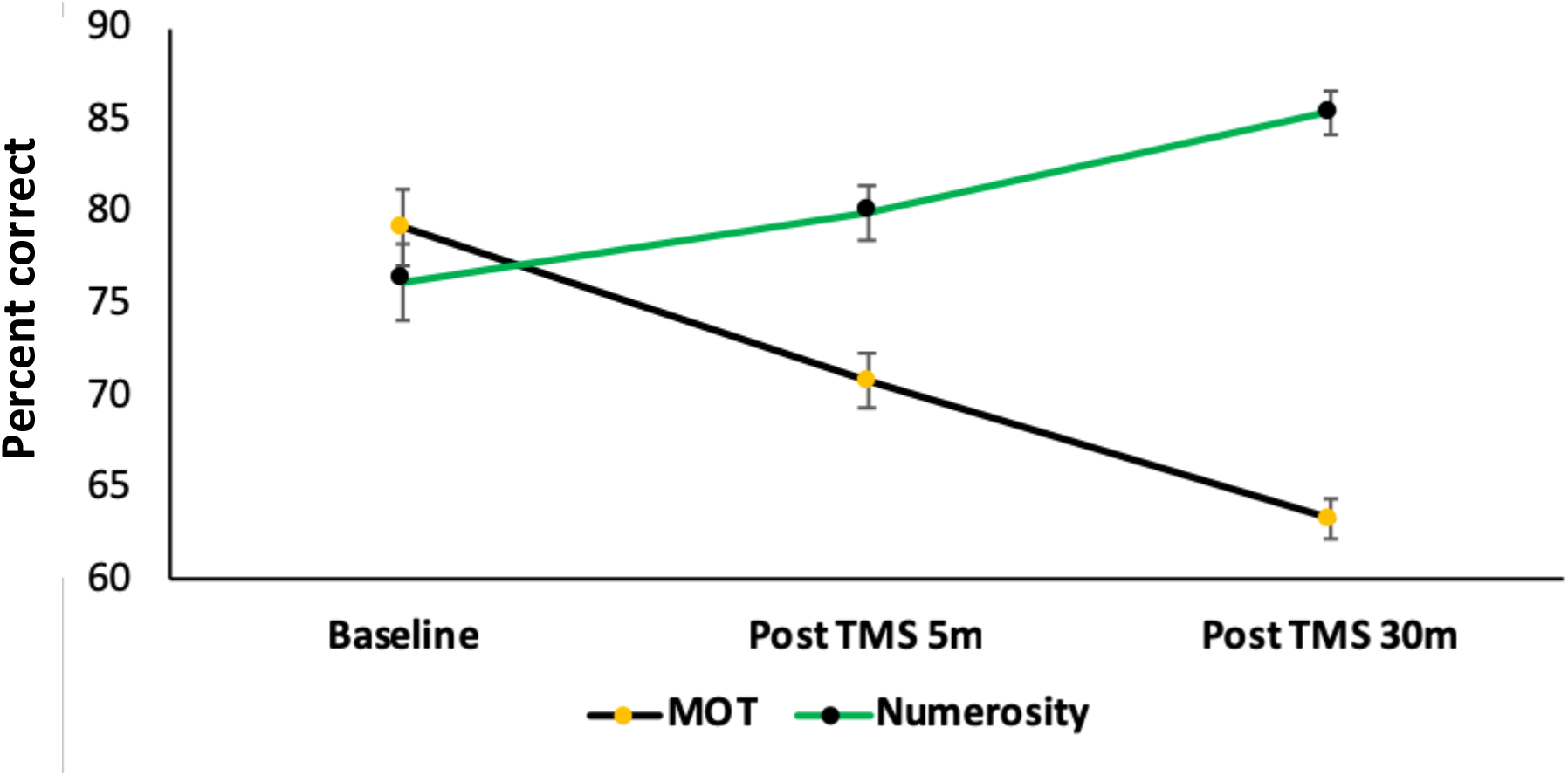
Percent correct MOT task accuracy at the stimulated (right) hemifield, following active cTBS, using multiple object tracking (black line) and numerosity judgement (green line) tasks. The blue, red and green lines indicate baseline, 5min post cTBS and 30-min post cTBS. Error bars show +/− 1 SEM.

## DISCUSSION

The aim of this study was to further assess the role of MT+ in attentive motion tracking. We found that left MT+ cTBS significantly impaired MOT task performance within the right visual hemifield at 5- and 30-mins post-stimulation. No impairment was observed for any of the control conditions (left visual hemifield for the active cTBS session and both hemifields for the sham cTBS session). In fact, MOT task accuracy improved 30 mins post-stimulation for all control conditions suggesting a task practice effect. The presence of a task practice effect suggests that the magnitude of the active cTBS-induced MOT accuracy reduction we observed may have been underestimated. The absence of any reduction in task performance with our control behavioural task (numerosity judgement) suggests that our results are not due to a non-specific effect of MT+ cTBS on psychophysical task performance.

Our results involving MOT are different from those reported by Battelli et al. (2009) who found that inhibitory 1hz rTMS of MT+ did not affect MOT task performance. This is most likely due to methodological differences between the two studies. Battelli et al., 2009 used 1 Hz rTMS, which was the gold standard at the time, as opposed to the cTBS protocol used in the current study. cTBS may have a more pronounced effect on cortical excitability than conventional rTMS (Cárdenas-Morales, Nowak, Kammer, Wolf, & Schönfeldt-Lecuona, 2010; Huang et al., 2005), although this has not been tested directly for area MT+. Moreover, in the Battelli et al., 2009 study, participants received both AIPS and MT+ stimulation in the same session. The large effect of IPS stimulation may have masked any effect of MT+ stimulation. Finally, because our study focused only on hMT+, we were able to introduce a within-session hemisphere control as well as a separate sham stimulation session. These controls may have allowed us to isolate the effect of hMT+ stimulation with greater sensitivity.

Our results are consistent with a number of previous studies demonstrating that cTBS can modulate MT+ activity and generate unilateral changes in global motion perception (Cai et al., 2014; Chen et al., 2016). We extend this previous work to reveal an effect of MT+ cTBS on a task that may engage higher-level attentive motion tracking mechanisms. The role of MT+ in MOT task performance could be explained by a number of mechanisms. Previous studies have observed that MT+ cTBS only impairs performance of motion tasks that involve signal/noise segregation (Cai et al., 2014; Chen et al., 2016). Specifically, MT+ cTBS had no effect on motion direction discrimination for a stimulus consisting only of signal dots, whereas performance was impaired for a stimulus with both signal and noise dots. cTBS of motion sensitive area V3A had the opposite effect. Non-human primate neurophysiology also indicates that MT plays a crucial role in signal/noise segregation of motion signals (Rudolph & Pasternak, 1999). Therefore, MT+ may support segregation of target dot motion from distractor dot motion during MOT task performance. However, unlike global motion coherence tasks, the target and distractor dots in MOT stimuli cannot be discriminated on the basis of their motion trajectories. Rather, attentional mechanisms are required to monitor which dots belong to which group throughout stimulus presentation. Our results suggest that MT+ is an important part of the network that supports target and distractor segregation during attentive motion tracking. This contention is consistent with the known effects of attentional modulation on MT+ activity (Beauchamp, Cox, & DeYoe, 1997; Berman & Colby, 2002; Chawla et al., 1999; Jody C Culham et al., 2001; O’Craven, Rosen, Kwong, Treisman, & Savoy, 1997; Rees, Frith, & Lavie, 1997; Treue & Maunsell, 1996) and the reciprocal connections between MT+ and brain areas which are highly specialized for attention processing such as the AIPS in humans (Howe et al., 2009; Treue & Maunsell, 1996) and the ventral intraparietal area in monkeys (Maunsell & van Essen, 1983).

More generally, our results provide new insights into MOT deficits that have been reported for individuals with amblyopia (Giaschi, Chapman, Meier, Narasimhan, & Regan, 2015; Ho et al., 2006; Secen et al., 2011) or neurodevelopmental disorders such as William’s syndrome (O’Hearn et al., 2010, 2005) and autism (Koldewyn et al., 2013). Although our results do not rule out deficits in higher-level attention areas in individuals with these conditions, they do demonstrate that abnormal MT+ activity would be sufficient to cause an attentive motion tracking impairment. In agreement with this conclusion, amblyopia is known to cause abnormal MT+ activity (Bonhomme et al., 2006; Ho & Giaschi, 2009; Thompson, Villeneuve, Casanova, & Hess, 2012) and the most robust reduction of BOLD response, associated with MOT deficit in amblyopia, was found in MT+, rather than other attentive motion processing areas, such as IPS and FEF (Secen et al., 2011). In addition, both William’s syndrome (Atkinson et al., 1997, 2003; Atkinson et al., 2006; Palomares & Shannon, 2013; Reiss et al., 2005) and autism (Brieber et al., 2010; Manning, Charman, & Pellicano, 2013; Spencer et al., 2000; Tsermentseli, O’Brien, & Spencer, 2008) have been linked to impaired global motion perception for noisy stimuli. This indicates an MT+ impairment that may be sufficient to explain any attentive motion tracking deficits. In summary, MT+ deficits may have a widespread effect on visual function that extends to attentive tracking of moving targets.

## Acknowledgements

This work was funded by NSERC grants RPIN-05394 and RGPAS-477166.

## CRediT author statement

**Arijit Chakraborty:** Conceptualization, study design, data collection, statistical analysis, writing original manuscript draft.
**Tiffany T. Tran:** Data collection, manuscript review. **Andrew E. Silva:** Data analysis, manuscript review. **Deborah Giaschi:** Conceptualization, study design, manuscript review and editing. **Benjamin Thompson:** Conceptualization, study design, manuscript review and editing, supervision

